# Mice use robust and common strategies to discriminate natural scenes

**DOI:** 10.1101/156653

**Authors:** Yiyi Yu, Riichiro Hira, Jeffrey N. Stirman, Waylin Yu, Ikuko T. Smith, Spencer L. Smith

## Abstract

Mice use vision to navigate and avoid predators in natural environments. However, the spatial resolution of mouse vision is poor compared to primates, and mice lack a fovea. Thus, it is unclear how well mice can discriminate ethologically relevant scenes. Here, we examined natural scene discrimination in mice using an automated touch-screen system. We estimated the discrimination difficulty using the computational metric structural similarity (SSIM), and constructed psychometric curves. However, the performance of each mouse was better predicted by the population mean than SSIM. This high inter-mouse agreement indicates that mice use common and robust strategies to discriminate natural scenes. We tested several other image metrics to find an alternative to SSIM for predicting discrimination performance. We found that a simple, primary visual cortex (V1)-inspired model predicted mouse performance with fidelity approaching the inter-mouse agreement. The model involved convolving the images with Gabor filters, and its performance varied with the orientation of the Gabor filter. This orientation dependence was driven by the stimuli, rather than an innate biological feature. Together, these results indicate that mice are adept at discriminating natural scenes, and their performance is well predicted by simple models of V1 processing.

## Introduction

Visual processing of natural scenes is essential for animal survival. Mammalian visual systems including primates and rodents evolved to efficiently process natural stimuli^1-5^. Mice use vision to hunt prey^6^, avoid danger^7-9^, and navigate^10,11^.

A number of studies have characterized the ability of mice to discriminate visual stimuli including gratings^12-14^, simple shapes^13-16^, and random dot kinematograms^17^. However, these results cannot be extrapolated to natural scene discrimination, because visual coding is substantially different between natural images and artificial ones^4,5,18^. Moreover, the spatial resolution of mouse vision is orders of magnitude lower than that of primates and carnivorans^12,19-23^. Even when natural scenes might be discriminated, individual mice could focus on different regions of the images to discriminate them, and this would lead to high mouse-to-mouse variability. Thus, investigating natural scene discrimination in mice can provide essential information for understanding evolved encoding strategies of mammalian visual systems.

The perception of visual information depends on processing by primary (V1) and higher visual cortical areas^24-28.^ One prominent feature of V1 neurons is their orientation tuning^29,30^. This selectivity can facilitate the sparse coding of natural images by V1 neurons^1-3^. Orientation specific features are further transformed and integrated in higher visual areas to extract higher order statistical structures of the image and detect objects^31-33^. Thus, the orientation selectivity is a foundation of visual perception. However, it is unclear how orientation features in naturalistic images can contribute to animal behavior.

Here, we developed a natural image discrimination task for freely moving mice using an automated touchscreen-based system. We found that mice successfully and quickly learned to discriminate images of natural scenes, the mouse-to-mouse consistency was high, and their performance could be well predicted by a simple model of V1 encoding.

## Results

### Mice learned to discriminate natural scenes

We used the automated touchscreen-based system^16,34^ that we previously adapted for visual discriminations^17^ (**Fig. 1a**). In the task, mice were presented with two images simultaneously, each in one of two presentation windows on the screen. The mice learned to touch a target image, avoiding a distractor image, to get a reward. Thus, it is a type of two-alternative, forced-choice (2AFC) task. All mice trained in the main experiment (6 of 6) successfully passed the pre-training phases (see **Methods**), meeting criteria to advance to natural image discrimination (NID) training in 14.5 ± 2.9 days (mean ± S.D.) (**Fig. 1b**). These mice also readily acquired the NID training task (6 of 6), ultimately discriminating correctly between a natural target image and 10 distractor images on 85% or more of trials (**Fig. 1c,d**). Two out of six mice were trained for an hour per day, and the other four mice were trained for two hours per day. Once mice performed the NID training task with 85% accuracy for two consecutive days, they moved to the NID testing phase. Mice required fewer training sessions to reach criterion for NID compared to the mice trained in the random dot kinematogram (RDK) task we previously reported^17^ (3, 3, 3, 5, 8, 9 days for NID vs. 5, 10, 11, 14, 15, 18 days for RDK; p = 0.0088, *t*-test).

**Fig. 1.**
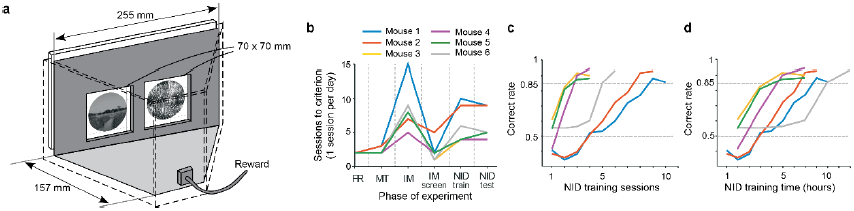
Touchscreen-based training system and learning curves. (a)Touchscreens displayed stimuli and registered responses in the behavior training system. Mice learned to touch the target image and avoid a distractor to get a reward. (b)Mice progressed through a series of pretraining phases within about a month (FR, Free Reward; MT, Must Touch; IM, Image; NID, Natural Image Discrimination; phases are explained in **Methods**). (**c**) Mice learned to perform the NID task to criterion (correct on 85% of trials) in 10 sessions or less, which corresponded to (**d**) 12 hours or less of training (data in panels c,d are the same, but are plotted against different units on the X-axis).

The NID testing phase consists of testing blocks and interleaved training blocks (**Fig. 2a**; see **Methods**), a strategy we used in our prior work^17^. In testing blocks, one of the 12 distractor images were selected for each trial. In the interleaved training blocks, the distractor image was always the same, and was easy to discriminate from the target. The target image was always the same in both block types. The mice had to touch the target image to receive a reward in the interleaved training blocks, otherwise they received a time-out. By contrast, mice received a reward on every trial, when touching either of the images in the testing block. Mice are never cued as to which trial type they are in. We find this approach effective^17^, perhaps because the always-rewarded testing blocks prevent the mice from getting too frustrated by the difficult discrimination trials, while the interleaved training blocks keep the mice honest. Five out of six mice performed correctly on > 85% of the interleaved training trials, which indicated that they were performing the task correctly, and they were included for further analysis. By contrast, one mouse (Mouse 6) had a lower correct rate for interleaved training blocks (**Supplementary fig.1**). This mouse seemed to recognize that it does not have to touch the target image to get rewards during the testing blocks, and it tended to select the left panel. Accordingly, we excluded this mouse, and analyzed the data in the testing blocks from the remaining five mice. We computed the correct trial rate with all five animals for the testing blocks (range: 1920 - 2712 trials, over 4 – 8 sessions per mouse). Repeated trials with the same sets of a target and distractor (range: 60-226 trails per image pair) enabled us to precisely estimate the correct trial rate.

**Fig. 2.**
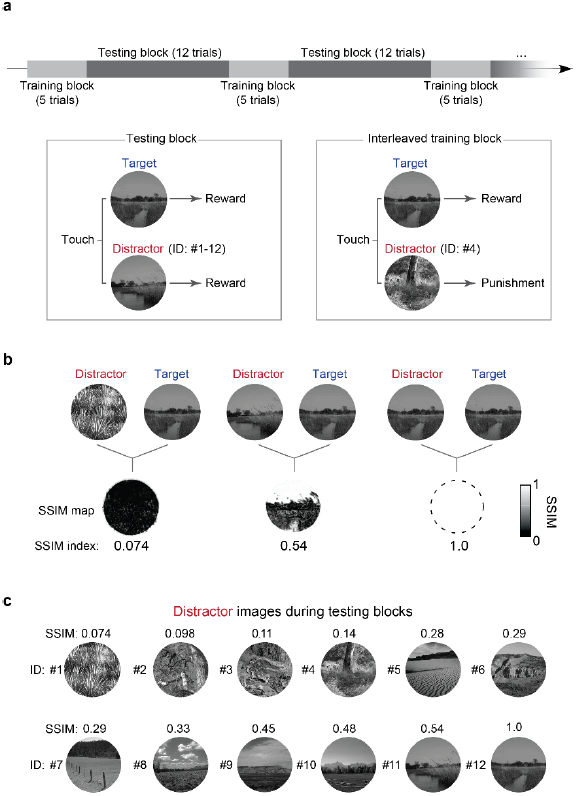
Testing procedure and SSIM index for quantifying expected difficulty. (**a**) NID testing sessions were composed of two types of blocks, testing blocks and interleaved training blocks. In testing block trials, one of 12 distractor images was presented with a target image. In interleaved training trials, the same distractor image was always presented with a target image. The incorrect choices were only punished (with a time-out and no reward) in the interleaved training trials. (**b**) Distractor and target images displayed in a trial were compared using structural similarity (SSIM). An SSIM map was obtained for each pair, and averaged yield a scalar SSIM index for the image pair. (**c**) Twelve distractor images were used, which varied in SSIM index with the target image (see also **Supplementary Table 1.**). One distractor (ID: 12) was the same as the target image, and served as an internal control.

### Behavior performance was predicted by structural similarity between images

To create psychometric curves to analyze mouse performance on this task, we need a metric that corresponds to the difficulty of each discrimination. For example, for discriminating gratings with different orientations, the orientation difference would be the appropriate metric to use. With natural images, the choice of metric is not straightforward, and multiple metrics could suffice. We chose to estimate the image similarities between two simultaneously presented images using the structural similarity (SSIM) index metric. The SSIM indices for all pairs of presented natural stimuli were calculated as reported by Wang et al. 2004 (Ref. ^35^) (see **Methods**). SSIM indices have been used to estimate the discriminability of artificial image pairs in prior mouse behavior studies^36,37^. In the testing blocks, the SSIM indices for image pairs ranged from 0.074 for the most dissimilar images, to 1 for trials when the same image was displayed on both sides of the screen (**Fig. 2b,c; Supplementary table 1,2.**). Psychometric curves were plotted using the correct trial rate and SSIM indices for 12 distractor images (**Fig. 3a**). The performances of the mice were remarkably similar (the thresholds of the psychometric curves were 0.29, 0.30, 0.27, 0.34 and 0.32; **Fig. 3b**). In total, the SSIM index approximated the correct trial rate, and the threshold was 0.30 ± 0.03 (mean ± S.D.) (**Fig. 3c**). To quantify the inter-mouse similarity, we computed the coefficient of variation (CV) for the psychometric threshold. The CV for the psychometric threshold in the natural scene discrimination task was 0.089, which is much smaller than the CV in the global motion discrimination task (0.24) that was carried out using the same apparatus^17^. Thus, the performance in the natural scene discrimination task is highly reproducible mouse-to-mouse.

**Fig. 3.**
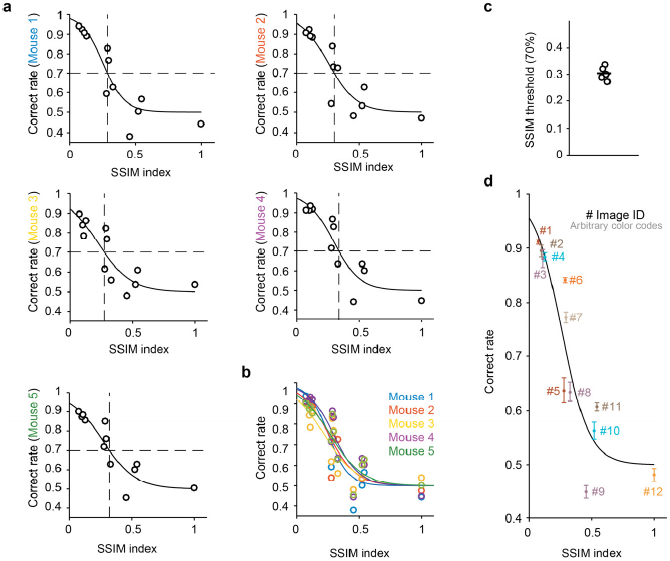
Psychometric curves for natural image discrimination. (**a**) During the testing blocks, mice were presented stimulus pairs of SSIM indices between 0.074 and 1 in random order, allowing us to sample the full range of SSIM indices for each animal. The same pairs were shown repeatedly to generate accurate estimates of the correct rate for each data point. Psychometric curves were fit to the data. The SSIM threshold (vertical dotted line; percent coherence at 70% accuracy) was computed from the fitted curves. (**b**) The psychometric curves from the five mice were similar. (**c**) The SSIM thresholds for the five mice were also very similar. (**d**) Each data point shows mean ± S.E.M. for each distractor image across all five mice. The psychometric curve is for the averaged data (threshold = 0.30). Note that for some distractors (e.g., #5, #9, and #11), the correct rate was not accurately approximated by the psychometric curve.

### High inter-mouse agreement

The SSIM index-based psychometric curves fit the data well, but behavioral data for some image pairs deviated from the psychometric curves (**Fig. 3d, 4a**). Notably, this deviation was not due to outlying data points from a few mice, but instead, data from all mice similarly deviated. In fact, the correct rates for each mouse were highly predictable by the mean correct rate of the other four animals (**Fig. 4b**). We analyzed the residuals of the fits by computing the root mean squared error (RMSE) between a predictor and the actual data. The RMSE for the simple case where the correct rate for each mouse was predicted by the mean correct rate of the other mice (mean ± S.E.M.: 0.047 ± 0.0046) were significantly smaller than the RMSE values for the SSIM-based psychometric curve fits (mean ± S.E.M.: 0.079 ± 0.0026) (**Fig. 4c**; p = 0.00053, paired *t*-test). These results indicate that the mouse visual system uses strategies that are not fully captured by SSIM indices.

**Fig. 4.**
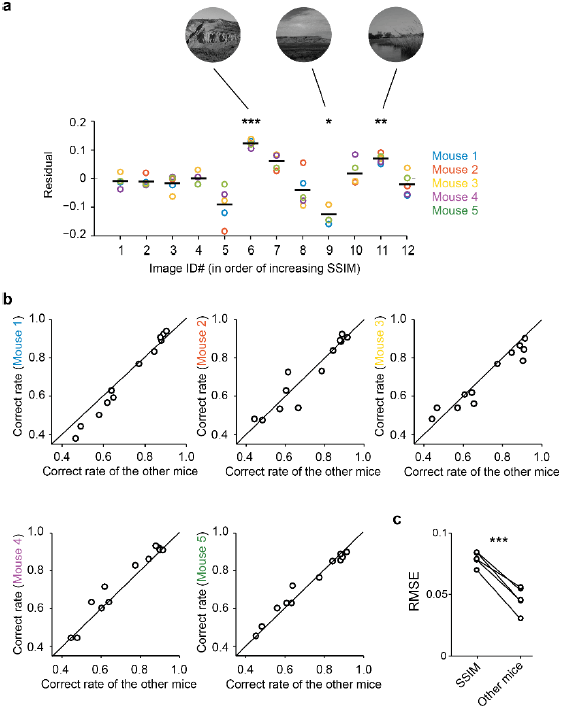
High inter-mouse agreement. (**a**) To examine the difference between the actual correct rate of each mouse and the predicted correct rate by the SSIM-based psychometric curve for each distractor image, we plotted the residuals. For some images, the mice collectively deviated from the predicted correct rate (ANOVA, F(48, 11) = 17.8, p= 2.3 x 10^-13^; *t*-test, ^*^p<0.05^, **^p<0.01, ^***^p<0.001). (**b**) The correct rates of each mouse were plotted as a function of mean correct rates of the other mice, for each distractor image (for reference, the unity line is shown in black). The correct rate of each mouse can be accurately predicted by that of other mice. (**c**) The RMSE of the inter-mouse agreement was smaller (i.e., more accurate) than that from SSIM-based psychometric curves (^***^p<0.001; paired *t*-test).

The mouse visual system does not have sufficient acuity with which to distinguish each pixel of the touch screen in this apparatus. One of the highest reported behaviorally-measured acuities in mice is 0.49 cycles per degree (Ref. ^12^), and this corresponds to 2.2 pixels if mice view the screen from just 10 mm. Thus, in this apparatus, mice are discriminating lower resolution representations of the two images. To investigate whether high spatial frequency information was biasing the SSIM-based estimator, we filtered the image with Gaussian filter with standard deviation from 1 to 112 pixels (**Fig. 5a, b**), and recomputed SSIM indices after Gaussian filtering (fltSSIM). The fltSSIM reached a minimal RMSE when the standard deviation was 4.6 pixels, but the improvement was small. This indicates that high spatial frequency information in the natural images is not responsible for more than a small amount of the RMSE when estimating behavior with SSIM.

**Fig. 5.**
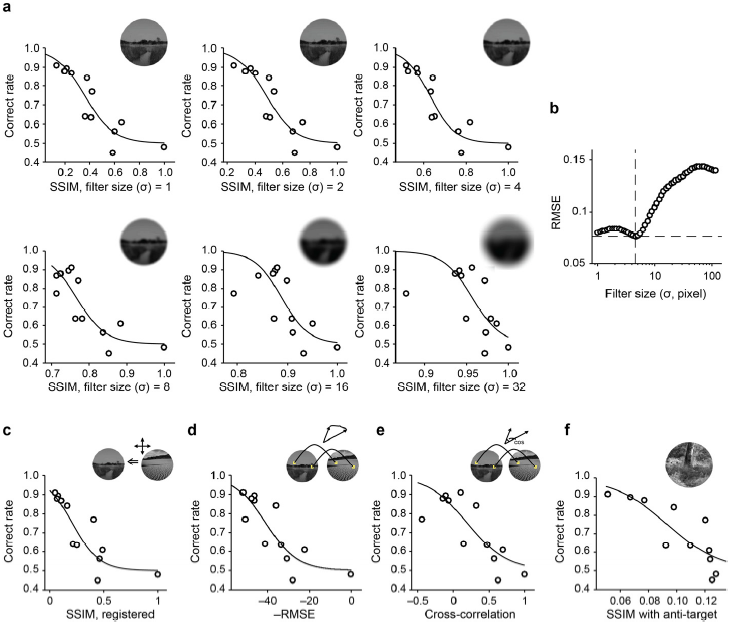
Blurring the images and other attempts to reduce the fit error (RMSE) Several approaches were investigated to determine whether they could reduce the error of the psychometric curve fit. (**a**) To determine whether high spatial frequency features of the images (which could be imperceivable to the mice, but still influence SSIM calculations) caused the relatively high root mean squared error (RMSE) values, the images were filtered (blurred) with Gaussian filters of a range of standard deviations (s). The inset shows the filtered target image. (**b**) Overall, blurring the images only marginally decreased the RMSE, and aggressive blurring led to even higher RMSE. (**c**) Registration of images prior to SSIM calculations did not improve the psychometric curve fit. (**d**) An alternative metric, pixel-wise RMSE between distractors and a target image, also did not improve the RMSE of a fit curve. (**e**) Another alternative metric, pixel-wise cross-correlation between distractors and the target image, also failed to yield low-RMSE curve fits. (**f**) Finally, calculating the SSIM between distractors and a distractor for the training phase (the “anti-target”), also failed to yield a low RMSE. RMSEs for (**c-f**) was 0.089, 0.095, 0.11 and 0.10, respectively.

SSIM is commonly used for estimating image similarity, but it is vulnerable to image translation. To robustly estimate the similarity between two images, we translated images so that the mean squared difference between two images was minimized. The SSIM values were then recomputed (SSIM after registration, regSSIM). However, the predictability of regSSIM was comparable to that of the original SSIM calculations, and RMSE still exceeded that of a simple inter-mouse predictor (**Fig. 5c**). Overall, SSIM analysis does not predict performance on this task as accurately as inter-mouse agreement.

SSIM is a sophisticated measurement with multiple components, and this sophistication could lead to biases that degrade performance as a predictor for mouse behavior on this task. Thus, we turned to two simple metrics to measure the similarity of two images: pixel-wise cross-correlation and RMSE between the two images. These parameters also failed to predict the correct rate as accurately as the mean performance of the other mice could (**Fig. 5d,e**). Finally, we tested the hypothesis that mice tended to avoid images similar to the distractor of the interleaved training phase (“anti-target”), because mice were only punished when they mistakenly selected interleaved anti-target images during the NID testing. However, recomputed SSIM indices using the anti-target did not yield an improved predictor (**Fig. 5f**). This is evidence against the hypothesis and indicates that the mice did not have tendency to avoid the “anti-target”. Overall, these results indicate that the mouse visual system uses strategies to judge the similarity between two images that are not captured well by SSIM or other metrics we examined. Therefore, we turned to a neurobiological model-based approach.

### V1-inspired model accurately predicts discrimination performance

So far, we have explored image comparison metrics that measure the similarity between two images. These metrics predicted mice behavior, but less accurately than a simple mean of other mice. Moreover, these metrics are not an intuitive model for the mouse visual system. Accordingly, we attempted to predict the discrimination performance with a model based on basic features of neuronal selectivity in mouse V1. The receptive fields of simple neurons in mouse V1 can be modeled as Gabor filters^38,39^. Each Gabor filter is convolved with a small patch of the image, and the result represents the degree of which that patch of the image and the Gabor filter are matched in orientation and wavelength (**Fig. 6a**). We convolved the target and distractor images with individual Gabor filters of various orientations and wavelengths, and calculated the similarity between the target and distractor images after filtering. We refer to this similarity as the orientation specific similarity (OSS) (**Fig. 6b**), since it is a function of the orientation of the Gabor filter (as well as wavelength). We plotted psychometric curves as the correct fraction versus OSS (**Fig. 6c**). We obtained RMSE values for these curves, for each set of wavelength and orientation parameters of the Gabor filters (**Fig. 6e,f**). The RMSE of the OSS model averaged over all orientations reached a minimal value when the wavelength of Gabor filter is approximately 7.07 pixels (**Fig. 6d**).

**Fig. 6.**
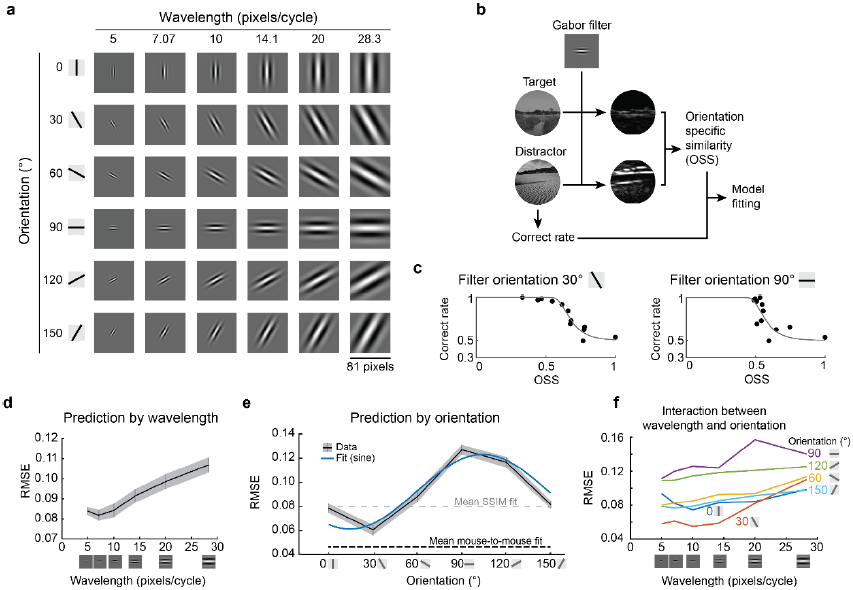
A simple Gabor filter model predicts mouse performance. (**a**) For a simple model the responses of V1 neurons, Gabor filters with various orientations and wavelengths were used to convolve the stimulus images. (**b**) The orientation specific similarity (OSS) between the target image and a distractor image was calculated by convolving a single Gabor filter with each image and then comparing the two results pixel-wise. The correct rate from the behavior data was then compared with these OSS values for each distractor image (compared to the target). (**c**) Example fits for the average correct rate across all mice against OSS values show that a filter with *θ* = 30°, *λ* = 10 (left) provided a good fit (RMSE = 0.055); and a filter with *θ* = 90°, *λ* = 10 (right) provided a poor fit (RMSE = 0.13). (**d**) Gabor filter patches with relatively short wavelengths provided lower RMSE fits to the behavior data (mean and S.E.M. are plotted; based on an average OSS over all orientations), and this suggests that relatively high spatial frequency information in the images is used by the mice for this discrimination. (**e**) The orientation of the Gabor patch used for calculating OSS influenced the resulting RMSE (p = 9 ×10^-11^, ANOVA; patches had wavelength = 10;mean and S.E.M. are plotted; blue curve is a sinusoidal fit). (**f**) An examination of the interaction between wavelength and orientation revealed that RMSE of the OSS-based fits depended more on orientation than wavelength of the Gabor filters.

The prediction RMSE of OSS varied by orientation (**Fig. 6e**, ANOVA p = 9.0 × 10^-11^). When OSS analysis used horizontally oriented Gabor filter, they performed poorly at predicting the correct trial rate, regardless of the spatial scale of the Gabor filters. By contrast, OSS analysis using a near vertically oriented Gabor filter predicted the animal behavior more accurately than the SSIM model. When OSS-based predictions were averaged over all orientations (using the optimal wavelength) the RMSE is similar to that of the SSIM model (using the optimal Gaussian filter, or blur) (RMSE of OSS, 0.082 ± 0.003 (λ = 7.07); RMSE of SSIM, 0.074 ± 0.003; *t*-test, p=0.08). The orientation bias in OSS-based prediction accuracy was consistent over a wide range of Gabor filter wavelengths (**Fig. 6f**). These results suggest that mice could use orientation specific features in the naturalistic image discrimination task.

The orientation specific prediction accuracy could result from an intrinsic bias of orientation selectivity in the mouse visual system^40,41^, or be acquired through the learning of specific images. To investigate these possibilities, we trained a new cohort of mice using the same set of images but rotated by 90° (n = 4 mice; **Fig. 7a**). If the orientation bias was intrinsic, then the orientation dependence of the OSS-based prediction should be identical to that obtained in the main experiment (**Fig. 6e**). If instead, the bias was learned based on the stimuli, then the orientation dependence of the OSS-based prediction should be rotated by 90°. All mice trained in the additional experiments (four of four) successfully passed the pre-training phases, just as the mice in the main experiments had. In the NID testing phase, we found that the orientation bias was shifted by ~90° (**Fig. 7b**). These results indicate that the orientation-dependence of the OSS-based prediction is not a result of a static innate orientation bias in the mouse visual system. Instead, the results suggest that the mouse visual system can learn to extract specific orientation information based on the natural images used in this behavior assay.

**Fig. 7.**
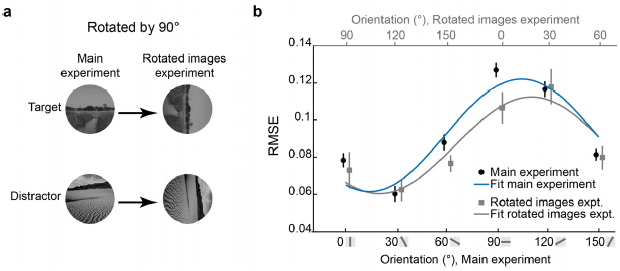
Experiments with rotated images showed that the orientation bias is not innate. (**a**) To examine whether the orientation-dependence of the OSS-based predictions was stimulus-based or innate, a separate cohort of mice was trained and tested on rotated versions of the same images. (**b**) The prediction error (RMSE) from the OSS-based model in the original experiment (black data points and blue sinusoidal curve fit; plotted against the bottom axis) was similar to the prediction error (RMSE) from the rotated images experiment (grey data points and grey curve; plotted against the top axis) after shifting the rotated image data by exactly 90 degrees (all OSS data used patches with wavelength = 10). Note that the bottom axis (for the main experiment) and the top axis (for the rotated images experiment) are offset by 90 degrees. This shows that the orientation specific prediction accuracy also shifted by 90 degrees.

## Discussion

Here we investigated the ability of mice to discriminate natural scenes, using an automated touchscreen-based system. We found that mice learned to discriminate natural scenes quickly, and that psychometric functions based on SSIM indices fit the data well. Further investigation revealed that the behavioral performance was highly consistent mouse-to-mouse, and deviated in significant and reproducible ways from the predictions from the SSIM-based psychometric curves. Thus, mice discriminate natural scenes using robust and common strategies, even when the images are displayed artificially, using an LCD monitor. To improve on the SSIM-based psychometric curves, we searched for a parameter or model that more accurately predicted the performance of the mice on this task. We found that a simple, V1-inspired model provides a prediction whose accuracy can approach that of the inter-mouse agreement. Thus, V1 processing may partly explain the way in which the mouse vision discriminates natural images.

Natural scenes were compared using the SSIM index, which is commonly used to estimate the similarity of two images^35^. SSIM indices correlated with the correct rate of the mice in our task. However, the psychometric curves plotted against SSIM provided only marginal fits to the data. In particular, there were several images that had significantly higher or lower correct rates across all mice than predicted by SSIM (**Fig. 4a**). In addition, the correct rates of mice were highly correlated, which indicated that the deviations from the SSIM-based psychometric curves were not due to mouse-to-mouse variance, but rather that the mice perceived the similarity of the images in ways not captured by SSIM.

Reducing the spatial resolution (i.e., blurring) of the images to better match mouse vision did not substantially improve the fits of SSIM-based psychometric functions, nor did several other approaches that we explored. Instead, we found that a Gabor filter-based approach provided the best fit. Notably, the improvement was specific to the orientation angle of the Gabor patch used. Horizontal orientation filtering did not provide a good fit, but a peri-vertical one did. This orientation bias of the Gabor filter-based approach was not due to innate properties of the mouse visual system. We know this because we tested a separate cohort of mice on the same images rotated by 90 degrees, and the orientation bias of the Gabor filter-based approach was rotated by the same amount. These results indicate that mice could extract orientation specific information depending on the task context.

In mammals, the orientation information itself could be modulated depending on the current demands. For instance, when humans attend to a specific orientation, there is increased activity in the portions of visual cortex that preferred the attended orientation^42^. In monkey V1, orientation-selective neurons fire at higher rates when attention is focused in the neurons’ receptive fields^43^. Also, the direction and orientation specificity of mouse V1 neurons increased during learning of a visual discrimination tasks when the preferred direction of the neuron was task relevant^44,45^. Thus, the orientation specific enhancement of visual processing appears to be a common feature of mammalian visual systems, and enhanced visual processing for specific orientations might explain the results from the present study. These changes in neural responses based on orientation could arise through learning.

Visual processing tasks can be categorized into scene/object recognition and motion/location recognition, which are represented along different “streams” or subnetworks of cortical visual areas^28,46,47^. In that context, this NID task complements the random dot kinematogram task we recently developed^17^. These experimental paradigms can be used to investigate stream-specific higher visual processing of mice^25,26,28,48.^

## Methods

### Subjects

Ten adult C57BL/6 mice (two males and four females in main experiments and two males and two females in additional experiments) were used in the experiments reported here. Animals were between 80 and 160 days old at the start of training, which lasted for approximately one month. Mice were housed in rooms on a reversed light-dark cycle (dark during the day, room light on at night), and training and testing were performed during the dark cycle of the mice. All training and procedures were reviewed and approved by the Institutional Animal Care and Use Committee of the University of North Carolina, and performed on campus, which is accredited by the Association for Assessment and Accreditation of Laboratory Animal Care (#329).

### Apparatus

The operant chamber and controlling devices were the same as previously described^17^, and are based on work by Saksida and Bussey^15,16,34^. Briefly, a touchscreen panel was on the long size of a trapezoidal chamber, opposite of liquid reward port (a strawberry-flavored yogurt-based drink; Kefir). Correct responses were indicated by a brief auditory tone, and the reward port was illuminated. During time-outs, a brief burst of white auditory noise was played, and the chamber light was illuminated for the duration of the time-out.

### Stimuli

For the NID task, we selected pictures of natural scenes (430 by 430 pixels static images, JPEG format). The original set of images was taken from three naturalistic image databases:UPenn(http://tofu.psych.upenn.edu/~upennidb/)^49^,McGill(http://tabby.vision.mcgill.ca/)^50^, and MIT (http://cvcl.mit.edu/database.htm)^51^. Average luminance of images was adjusted to be similar. One image was used as the target image during NID training sessions and NID testing sessions. Ten images that have low SSIM with the target image, and eleven images with varied SSIM (plus one image that is the same as the target image) were selected as distractor images on the NID training and NID testing sessions, respectively (**Supplementary table 1,2)**. All images were masked by a circular aperture. During NID testing sessions, 5 interleaved training trials were provided after every 12 testing trials. One image with low SSIM (of the eleven distractors) was used as the distractor for the interleaved training.

### Behavioral Training

Food restriction and training stages 1-3 (FR, free reward; MT, must touch; IM image discrimination) for the 2AFC task were conducted as previously described^17^. Briefly, during the FR phase (training stage 1), mice learned to lick the reward spout to receive a liquid reward. As a result, mice associated the tone with the delivery of a reward, and learned the location of the reward. During the MT phase (training stage 2), mice had to touch any location on the screen at the front of the box to receive a reward. The goal of this phase was to associate touching the screen with delivery of a reward. During the IM phase (training stage 3), a simple black-and-white dot and fan image pair stimulus^16^ was presented, and mice were required to touch a specific target stimulus on the screen to earn a reward. On the NID training phase (training stage 4), mice had to touch the target image, avoiding one of 10 distractors. SSIM indices of all pairs were less than 0.2 (low SSIM implies that the two images are not similar). This stage of training incorporated correction trials as described previously^17^.

### Behavioral Testing

The NID testing condition was similar to the testing condition for kinematogram discriminations we previously described^17^. Images for the testing phase consisted of one target image and 12 distractor images whose SSIM indices ranged from 0.074 to 1. Testing sessions consisted of interleaved blocks of testing and training. Mice were not cued as to whether they were in a testing or training block. During testing blocks, all 12 distractors (12 trials per block) were presented in a random order, and all answers were rewarded. During training blocks (5 trials per block), stimuli with SSIM = 0.14 (i.e., very dissimilar to the target) were presented, and normal performance feedback (including time-out periods and correction trials) were provided. Performance during these interleaved training blocks served as an internal control to ensure the mice were still working to distinguish the stimuli in the testing blocks. Testing data was analyzed only if the animal performed on average at criteria (≥ 85% correct) during the interleaved training blocks. This criterion excluded one mouse out of the six mice in the main experiment. Testing sessions were conducted for 120 - 150 minutes per day. In additional experiments, the target and distractor images were rotated only in the NID training and testing sessions. All four mice generated testing data in the additional experiments.

### SSIM (Structural similarity)

SSIM indices for all pairs of presented natural stimuli were calculated as reported by Wang et al 2004 (Ref. ^35^). First, the pair of two images were smoothed with Gaussian filter (*σ*= 1.5 pixels). SSIM index was obtained for each window of a pair of images. First, the SSIM for each pixel was calculated for square windows, centered at the same pixel (*w,h*) of two images. The length of a side of the square window was eight pixels. The SSIM (*x,y*) was then obtained for two windows in a target image (*x*) and a distractor image (*y)* as follows,

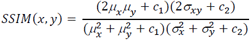

where *μ*_*x*_*, μ*_*y*_ are pixel intensity averages of window *x* and *y*, *σ*_*x*_, *σ*_*y*_, *σ*_*xy*_ are standard deviation of window *x* and *y*, and covariance of two windows, *c*_*1*_ and *c*_*2*_ are (0.01 x *L*)2 and (0.03 x *L*)^2^, respectively, where *L* is 255 (corresponding to the dynamic range of 8-bit monochrome images).

SSIM(*x,y*) was obtained for all (*w,h*) and was averaged over the circular area where the natural images are located. The average of SSIM(*x,y*) was called simply the “SSIM index” for each pair of target and distractor images in this paper (**Fig.2b**). One distractor was the same as a target image, so that its SSIM index = 1. The other image pairs had SSIM indices < 1.

### Psychometric curve

Psychometric curves were obtained with a regression of the following function (Weibull function) to the data,

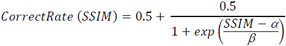

where *α* and *β* are parameters determined by the regression to each data set. The threshold was taken as the SSIM value (from this function) that corresponded to 70%accuracy.

### Image processing

Pixel-wise correlation and root mean squared error (RMSE) between a target image and each distractor image were obtained as candidate parameters that may capture the similarity of two images. The registration of two images were done with a MSE based registration algorithm (Turboreg)^52^.

### Gabor filter

We used Gabor filtering to analyze the orientation specific features of the images:

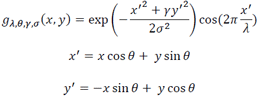

The Gabor filter bank was generated using the built-in Matlab function “gabor”. The wavelength (*λ*) of the Gabor filters ranged from 5 – 28 pixels per cycle. The orientation (*θ*) of Gabor filters was either 0°, 30°, 60°, 90°, 120°, or 150°. We set the aspect ratio (*γ*) of all Gabor filters to 0.5. The standard deviation of the Gaussian envelops, σ = 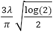 determined by the wavelength. We computed the Gabor feature magnitude *G*_*A,0*_*(x, y)* by convolving the Gabor filter (with a specific orientation and wavelength) with each image. Then we computed the orientation specific similarity (OSS) between two images by the following function:

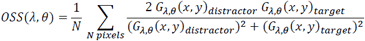

### Statistics

Student’s *t*-test was used for statistical comparisons. Pairwise comparisons were two-tailed. Error bars in graphs represent the mean ±S.E.M. unless otherwise noted. Analysis of variance (ANOVA) was used for multiple comparisons, followed by *t*-test (presented p-values for *t*-test were Bonferroni corrected). No statistical tests were performed to predetermine sample size. No blinding or randomization were performed.

## Acknowledgements

This work was supported by grants to S.L.S. from the Human Frontier Science Program, NIH (R01NS091335 and R01EY024294), the Simons Foundation (SCGB 325407SS), and the Whitehall Foundation. Y.Y. was supported by a Helen Lyng White fellowship, R.H. was supported by fellowships from Naitoh and the Japan Society for the Promotion of Science, and J.N.S. was support by a Career Award from Burroughs-Wellcome.

---

**Supplementary Figure S1.**
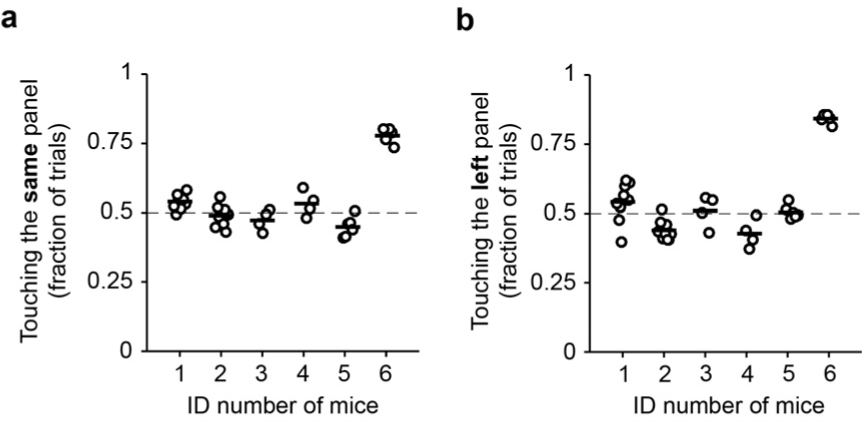
History dependency and choice bias. (**a**) Mouse number 6 tended to select the same side of the touchscreen on subsequent trials, while other mice did not. (**b**) Mouse number 6 tended to select the left panel, while the other mice were more even in their selections.

**Supplementary Table S1.**
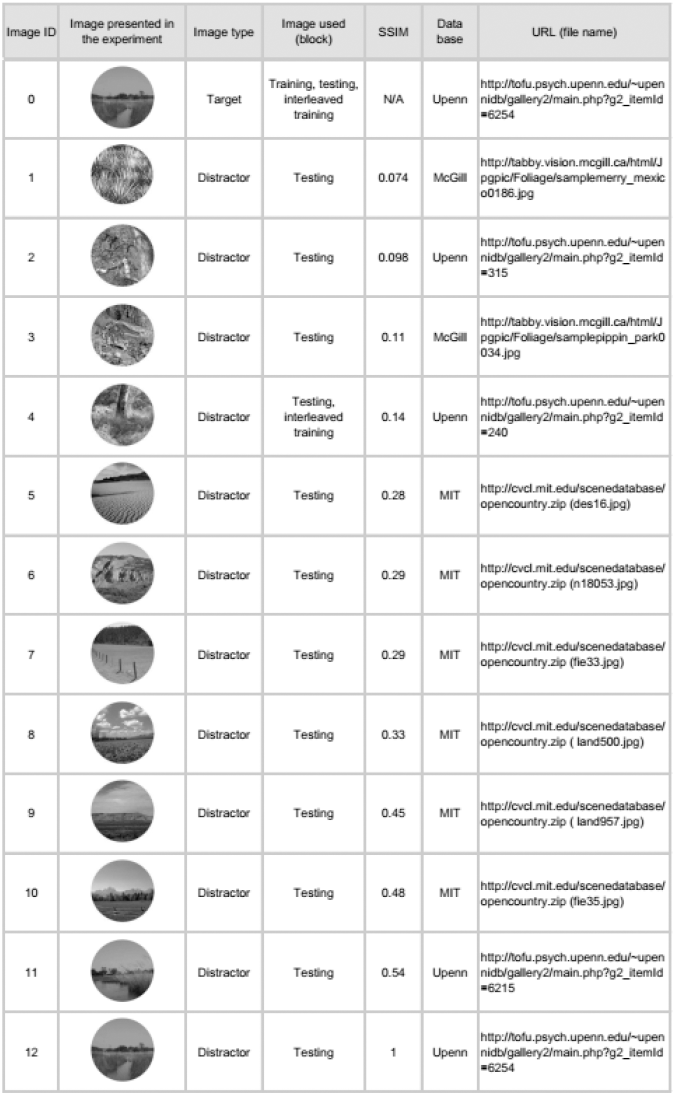
Images used during NID testing phase. The target image and all distractor images for NID testing block and interleaved training block are listed. The SSIM index and information for the database for each image are also listed.

| Image ID | Image presented in the experiment | Image type | Image used (block) | SSIM | Data base | URL (file name) |
| --- | --- | --- | --- | --- | --- | --- |
| 0 |  | Target | Training, testing, interleaved training | N/A | Upenn | http://tofu.psych.upenn.edu/~upennidb/gallery2/main.php?g2_itemId=6254 |
| 1 |  | Distractor | Testing | 0.074 | McGill | http://tabby.vision.mcgill.ca/html/Jpgpic/Foliage/samplemerry_mexico0186.jpg |
| 2 |  | Distractor | Testing | 0.098 | Upenn | http://tofu.psych.upenn.edu/~upennidb/gallery2/main.php?g2_itemId=315 |
| 3 |  | Distractor | Testing | 0.11 | McGill | http://tabby.vision.mcgill.ca/html/Jpgpic/Foliage/samplepippin_park0034.jpg |
| 4 |  | Distractor | Testing, interleaved training | 0.14 | Upenn | http://tofu.psych.upenn.edu/~upennidb/gallery2/main.php?g2_itemId=240 |
| 5 |  | Distractor | Testing | 0.28 | MIT | [http://cvcl.mit.edu/scenedatabase/opencountry.zip (des16.jpg)](http://cvcl.mit.edu/scenedatabase/opencountry.zip%28des16.jpg%29) |
| 6 |  | Distractor | Testing | 0.29 | MIT | [http://cvcl.mit.edu/scenedatabase/opencountry.zip (n18053.jpg)](http://cvcl.mit.edu/scenedatabase/opencountry.zip%28n18053.jpg%29) |
| 7 |  | Distractor | Testing | 0.29 | MIT | [http://cvcl.mit.edu/scenedatabase/opencountry.zip (fie33.jpg)](http://cvcl.mit.edu/scenedatabase/opencountry.zip%28fie33.jpg%29) |
| 8 |  | Distractor | Testing | 0.33 | MIT | [http://cvcl.mit.edu/scenedatabase/opencountry.zip (land500.jpg)](http://cvcl.mit.edu/scenedatabase/opencountry.zip%28land500.jpg%29) |
| 9 |  | Distractor | Testing | 0.45 | MIT | [http://cvcl.mit.edu/scenedatabase/opencountry.zip (land957.jpg)](http://cvcl.mit.edu/scenedatabase/opencountry.zip%28land957.jpg%29) |
| 10 |  | Distractor | Testing | 0.48 | MIT | [http://cvcl.mit.edu/scenedatabase/opencountry.zip (fie35.jpg)](http://cvcl.mit.edu/scenedatabase/opencountry.zip%28fie35.jpg%29) |
| 11 |  | Distractor | Testing | 0.54 | Upenn | http://tofu.psych.upenn.edu/~upennidb/gallery2/main.php?g2_itemId=6215 |
| 12 |  | Distractor | Testing | 1 | Upenn | http://tofu.psych.upenn.edu/~upennidb/gallery2/main.php?g2_itemId=6254 |

**Supplementary Table S2.**
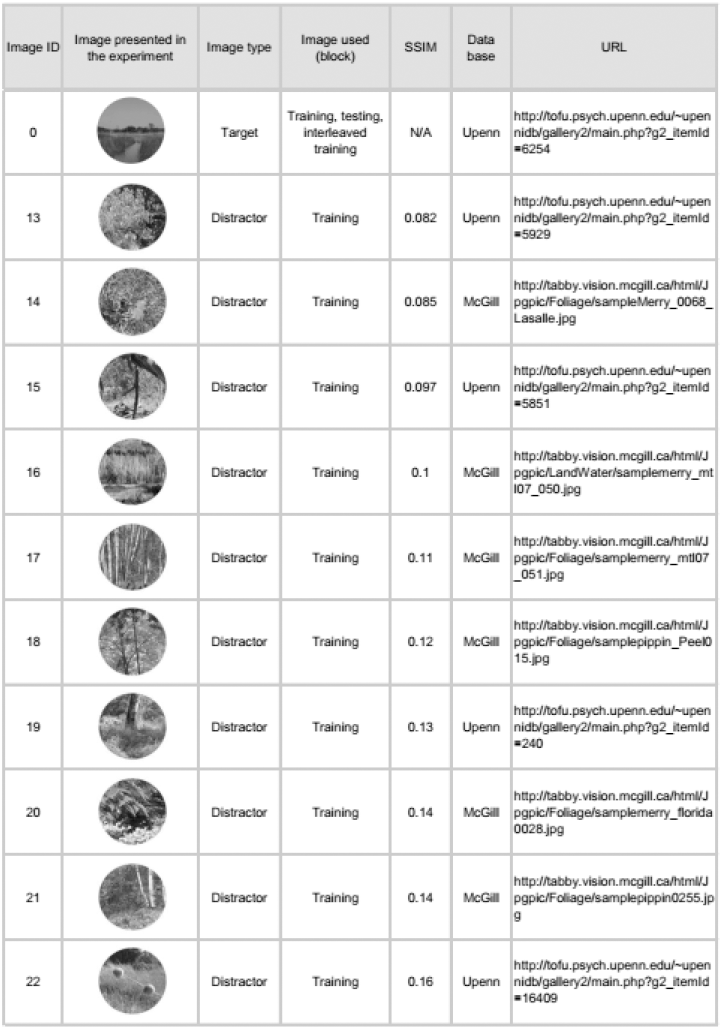
Images used during NID training phase. A target image and all distractor images for training phase are listed. The SSIM index and information for the database for each image are also listed.

